# Genetic admixture patterns in Argentinian Patagonia

**DOI:** 10.1101/586610

**Authors:** María Laura Parolin, Ulises F Toscanini, Irina F Velázquez, Cintia Llull, Gabriela L Berardi, Alfredo Holley, Camila Tamburrini, Sergio Avena, Francisco R Carnese, José L Lanata, Noela Sánchez Carnero, Lucas F Arce, Néstor G Basso, Rui Pereira, Leonor Gusmão

## Abstract

As for other Latin American populations, Argentinians are the result of the admixture amongst different continental groups, mainly from America and Europe, and to a lesser extent from Sub-Saharan Africa. However, it is known that the admixture processes did not occur homogeneously throughout the country. Therefore, considering the importance for anthropological, medical and forensic researches, this study aimed to investigate the population genetic structure of the Argentinian Patagonia, through the analysis of 46 ancestry informative markers, in 433 individuals from five different localities. Overall, in the Patagonian sample, the average individual ancestry was estimated as 35.8% Native American (95% CI: 32.2-39.4%), 62.1% European (58.5-65.7%) and 2.1% African (1.7-2.4%). Comparing the five localities studied, statistically significant differences were observed for the Native American and European contributions, but not for the African ancestry. The admixture results combined with the genealogical information revealed intra-regional variations that are consistent with the different geographic origin of the participants and their ancestors. As expected, a high European ancestry was observed for donors with four grandparents born in Europe (96.8%) or in the Central region of Argentina (85%). In contrast, the Native American ancestry increased when the four grandparents were born in the North (71%) or in the South (61.9%) regions of the country, or even in Chile (60.5%). In summary, our results showed that differences on continental ancestry contribution have different origins in each region in Patagonia, and even in each locality, highlighting the importance of knowing the origin of the participants and their ancestors for the correct interpretation and contextualization of the genetic information.

## Introduction

Argentina, like many other Latin American countries, has a large proportion of inhabitants with admixed ancestry, with a high Native American and European contribution and to a lesser extent from Africa. The most significant European migration occurred between 1860 and 1950. In that period, Argentina welcomed an estimated 5.5 million migrants, mainly from Spain and Italy [1,2]. According to the 1869 census, the Argentinian population had only 1.8 million inhabitants (National Institute of Statistics and Census of Argentina (INDEC), 2010), placing this migratory event as the most significant worldwide, considering the size of the recipient population. In this context, and for more than a century, the popular imaginary defined Argentina as a “white” country where most of its population was supposed to descend from European immigrants. However, the geographic distribution of the Europeans was strongly biased towards the city of Buenos Aires and the rest of the Central region of Argentina [1,2]. On the other hand, the Argentinian Patagonia was, until the last third of the 19th century, a virtually control-free area of the Republican state, a situation that allowed the native populations to preserve their autonomy for a long period [3]. This region lacked permanent settlings until the late 1800, when the so-called “Conquista del Desierto” (Desert Conquest) annexed the Patagonia to the Argentine Republic.

Until then, the Argentinian Patagonia had been inhabited mainly by hunter-gatherer native groups of low demographic density and high residential mobility. Although these groups had a long history of sporadic contacts with foreigners, only at the end of the 19th century there was a significant demographic change in the region, due to strong migratory flows coming from inland provinces and from bordering countries, mainly attracted by new labor opportunities [4].

Our research group has previously investigated the genetic composition of urban Argentinian populations with focus on the Patagonia region by means of uniparental and autosomal markers. In five populations of Chubut and Río Negro provinces, the analysis of mitochondrial DNA and Y-chromosome markers revealed heterogeneity through autochthonous and allochthone contributions. In this sense, mitochondrial analysis showed a higher Native American maternal contribution in populations of Esquel, San Carlos de Bariloche, Comodoro Rivadavia, Puerto Madryn and Trelew [3,5–8]. For the same populations, the Native American fraction of parental lineages, analyzed by Y-STR and Y-SNPs, was relatively high in the Andean populations of Esquel and San Carlos de Bariloche, and lower in the coastal cities of Comodoro Rivadavia, Puerto Madryn and Trelew [9]. These studies showed that a biased admixture process between European males and Native women lead to a partial replacement of native paternal lineages with allochthone ones. This process has been widely observed in other regions of Argentina [10–13], as well as in other countries in South America [14].

Regarding autosomal markers, the studies carried out in urban populations of Argentinian Patagonia varied on the number of populations, the source populations and the number of samples, and the genetic markers analyzed. In this respect, Toscanini et al. [15] analyzed 46 ancestry informative markers (AIMs) in samples from Santa Cruz and Neuquén provinces, and Corach et al. [12] studied 24 AIMs in samples from Rio Negro. These studies showed similar results for the Native American (32.3%-27.7%), European (68.4%-65.6%) and African (3.8%-2%) genetic contributions.

Studies carried out by our research team in Argentinian Patagonia comprised the analysis of blood systems [3,5,6], autosomal STRs [16], AIMs [17] and Alu markers [18]. Admixture results obtained showed relatively wide ranges of continental ancestry between markers: 44-29% Native American, 67-51% European, and 4-2% African. The study based on Alu markers, showed that the admixture process was not homogeneous across Patagonia region [18].

The most comprehensive ancestry studies by Seldin et al. [19], Wang et al. [20] and Muzzio et al. [21] were mainly focused on the differential distribution of Native American, European and African components in the Central and Northern regions of Argentina, while samples from the Southern region of the country were not analyzed.

The present study aimed to analyze a panel of 46 ancestry informative insertion-deletion markers (AIM-Indels), described by Pereira et al. [22], in a relatively large number of individuals (n = 433), to contribute to a better understanding of the genetic structure and different admixture processes that occurred in urban Argentinian Patagonia populations. For a more in-depth interpretation, with focus in the regional and local differences, the ancestry results were analyzed taking into account genealogical and historical data. This information allowed for investigating possible associations between genetic ancestries inferred by the 46 AIMs panel and the geographic origin of the participants.

The knowledge of the genetic composition of Argentinian Patagonia, in terms of individual and population ancestry composition will provide important information for medical, forensic and anthropological studies in the region.

## Materials and Methods

### Ethics Statement

This study was approved by the Ethics Committees of Puerto Madryn Zonal Hospital (Resol.009/2015) and San Carlos de Bariloche Zonal Hospital (Resol. 1510/2015). Moreover, it has been endorsed by Hospitals’ Teaching and Research Committees of Trelew, Comodoro Rivadavia and Esquel. Biological samples involved in the study were coded and anonymized, as per the Argentina National Law 25.326 of Protection of Personal Data. All samples were collected under written informed consent to participate in this study.

### Biological Samples and DNA extraction

A volume of 5 ml of blood was EDTA collected from 433 healthy non-related voluntary blood donors, randomly selected from hemotherapy services of public and private hospitals of five Argentinian Patagonia cities; two from the Andes region, namely San Carlos de Bariloche and Esquel, and three located in the coastal area: Puerto Madryn, Trelew, and Comodoro Rivadavia (Fig 1).

**Fig 1.**
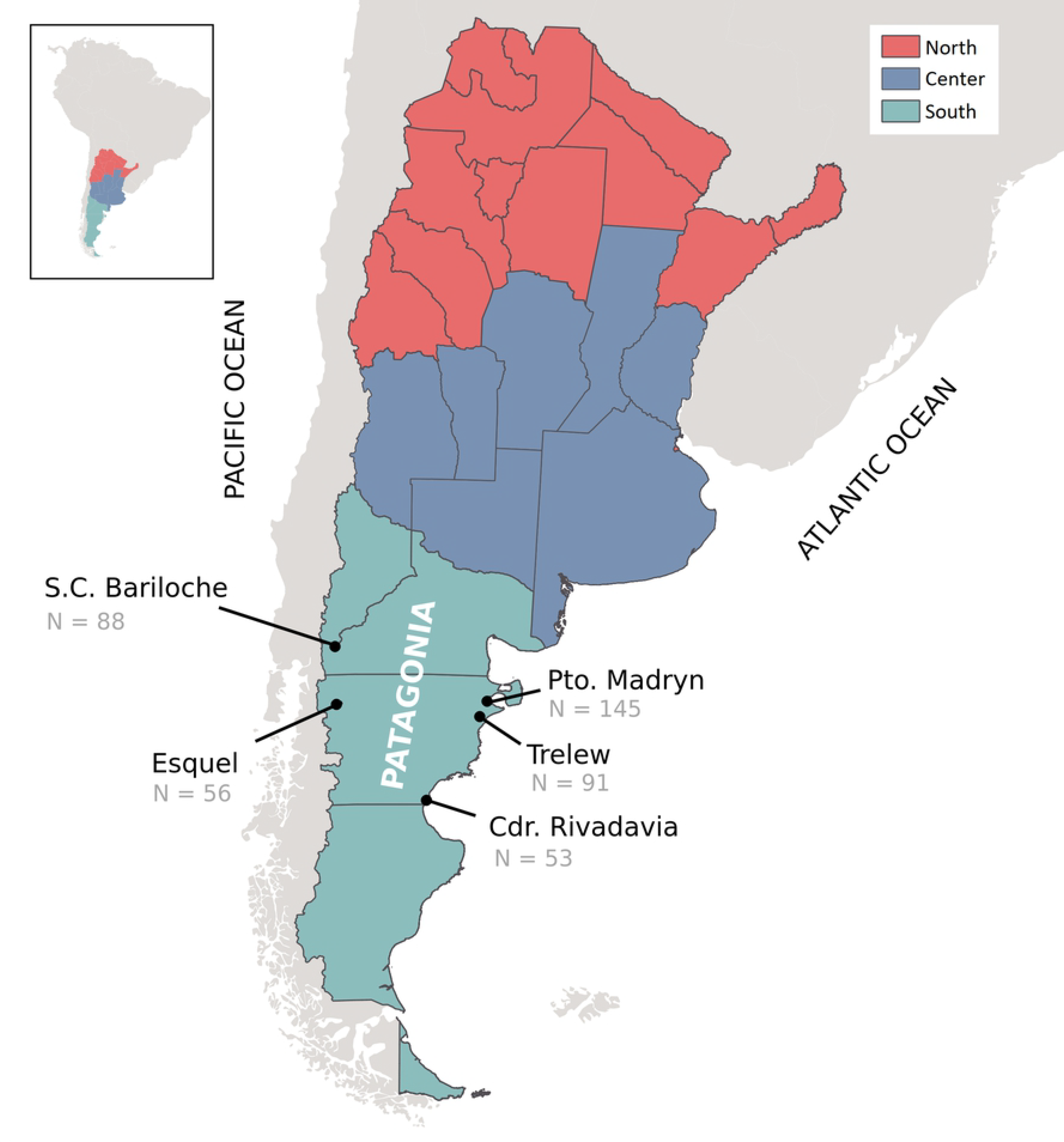
Map of Argentina showing the three main regions of the country, and the five studied populations.

Participants were thoroughly informed of the scopes of the study and answered a questionnaire to obtain information about their birthplace and that of the two preceding generations (parents and grandparents). This information allowed us to evaluate a possible association between genetic ancestry and geographic origin. DNA extraction was performed as described in Parolin et al. [18].

### Genotyping of the AIM-Indels

A panel of 46 AIM-Indels, developed by Pereira et al. [22] was genotyped in a single multiplex PCR followed by capillary electrophoresis. PCR thermocycling conditions were: an initial step of 15 min at 95°C; followed by 27 cycles at 94°C for 30 s, 60°C for 1.5 min, 72°C for 45 s; and a final extension at 72°C for 60 min. Dye-labeled amplified fragments were separated and detected using an ABI3130 Genetic Analyzer (Life Technologies), and automated allele calls were obtained with GeneMapper v.3.2 (Life Technologies). This panel showed to be highly informative to analyze genetic admixture in different Latin-American populations, including Argentina [15], Brazil [23], Bolivia [24] and Colombia [25,26].

### Statistical Analysis

Allele frequency estimations, Hardy-Weinberg exact tests, and population differentiation tests were performed using the Arlequin software v.3.5.2 [27]. Population genetic analyses were carried out comparing our five populations with others six Argentinian populations [15] and six South American populations from Brazil [23], Bolivia [24] and Colombia [25,26]. Genetic distances (*F*_ST_) between populations across 46 AIM-Indel markers were visualized in a multi-dimensional scaling (MDS) plot performed with the R v3.1.1 FactoMineR [28] statistical package. The proportion of genetic continental ancestry was estimated using the STRUCTURE v2.3.3 software [29]. In agreement with the historical and demographic settlement of Argentina we assumed a tri-hybrid model with three parental groups: Native American (NAM), European (EU) and African (AF) (i.e. K = 3). The allele frequencies of the Native American, European and African parental populations, from the diversity panel of human cell line samples HGDP-CEPH (Human Genome Diversity Project), were taken from Pereira et al. [22]. STRUCTURE runs consisted of 100,000 burning steps followed by 100,000 Markov Chain Monte Carlo (MCMC) iterations. For the Admixture model, the option "Use population Information to test for migrants" was used. Allele frequencies were correlated and updated only with individuals with POPFLAG = 1. Population admixture structure was represented in a principal component analysis (PCA), including individual genetic ancestry estimated in the samples studied and in the HGDP-CEPH parental populations. PCA plot was performed by the Stars package of R v3.1.1 [30]. Average individual ancestry, with a 95% confidence interval (CI), was estimated based on: i) parametric t distribution, if the sample size was higher than 29, and ii) non-parametric bootstrapping method, if the sample size was smaller than 30. Differences in the mean Native American, European and African ancestries, between the five studied populations and between the samples with four grandparents born in different geographic places, were analyzed using the non-parametric one-way ANOVA Kruskal-Wallis test [31]. This Non-parametric approach was selected because the distribution of the residuals of the parametric ANOVA deviated from normality. When the Kruskal-Wallis test was significant (p < 0.05), population pairwise differences were tested by the Dunn post-hoc analysis [32]. A Chi-squared test [33] was preformed to analyze the independence between the mean genetic ancestry obtained (Native American/European/African) and the regional origin (North/Center/South Argentina). Statistical analyses were performed by standard R v3.1.1 functions [30] implementing *ad-hoc* scripts. Geographic representations of the birthplace and ancestry proportions of the donors were obtained by ordinary kriging using the Geostatistical Analyst module of ArcMap 9.3 (ESRI). The covariances of the studied parameters (percentage of ancestry) were modeled in terms of the distance between sampling units [34].

## Results and Discussion

The genotyping results obtained in the five urban populations from Argentinian Patagonia are shown in S1 Table. Allele frequencies and expected heterozygosities were estimated for the 46 AIM-Indels are presented in S2 and S3 Tables, respectively. Exact test showed no significant deviations to the Hardy-Weinberg equilibrium expectations for all loci in the five populations (*p* ≥ 0.0038; for a significant level of 0.001, when applying the Bonferroni’s correction for 46 tests performed) [35].

### Genetic ancestry analysis

The estimation of continental admixture in 433 Argentine Patagonian individuals varied from 0.4 to 99.4% Native American, 0.3 to 99.2% European, and 0.2 to 23.4% African. Individual genetic ancestry obtained and its distribution through the HGDP-CEPH parental populations was represented in a Principal Component Analysis (PCA) (Fig 2).

**Fig 2.**
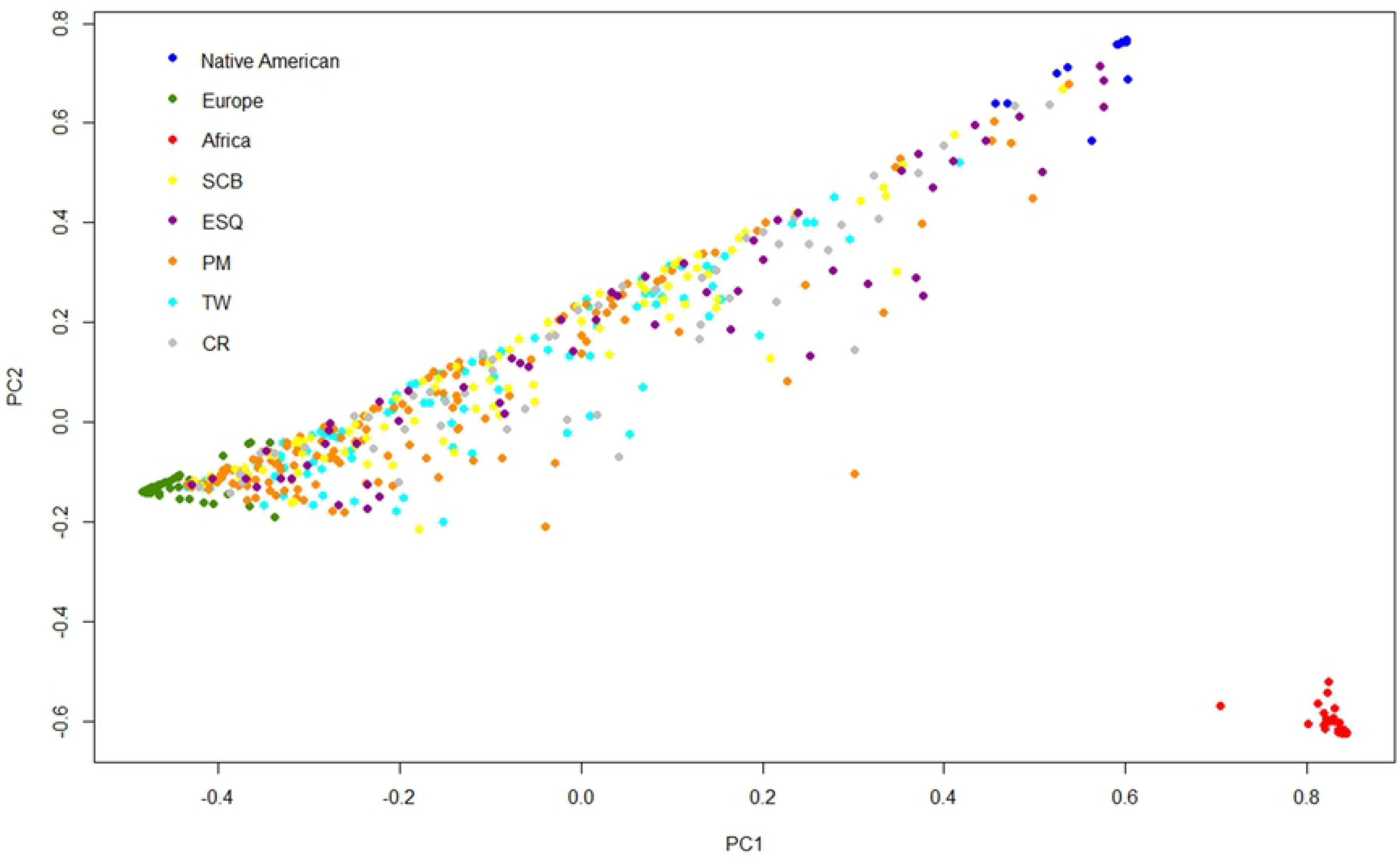
Principal Component Analysis plot showing the distribution of the Patagonian samples toward to the three HGDP-CEPH parental populations. SCB: San Carlos de Bariloche, ESQ: Esquel, PM: Puerto Madryn, TW: Trelew, CR: Comodoro Rivadavia.

The PCA plot showed a linear distribution of the Patagonia samples between European and Native American parental populations, with more density towards the European component, mainly for Puerto Madryn samples.

The average individual ancestry estimated in the total Patagonia sample was 35.8% (95% CI: 32.2-39.4%) Native American, 62.1% (95% CI: 58.5-65.7%) European, and 2.1% (95% CI: 1.7-2.4%) African. Comparing the five studied localities (Fig 3), a prevalence of the Native American ancestry was observed in Esquel and in Comodoro Rivadavia, located in western and southeastern Chubut, respectively. Meanwhile Puerto Madryn, from northeastern Chubut, was the Patagonian population with higher European contribution. San Carlos de Bariloche and Trelew were closer to the Native American and European averages in the global sample. The African ancestry was observed in similar low proportions in all populations.

**Fig 3.**
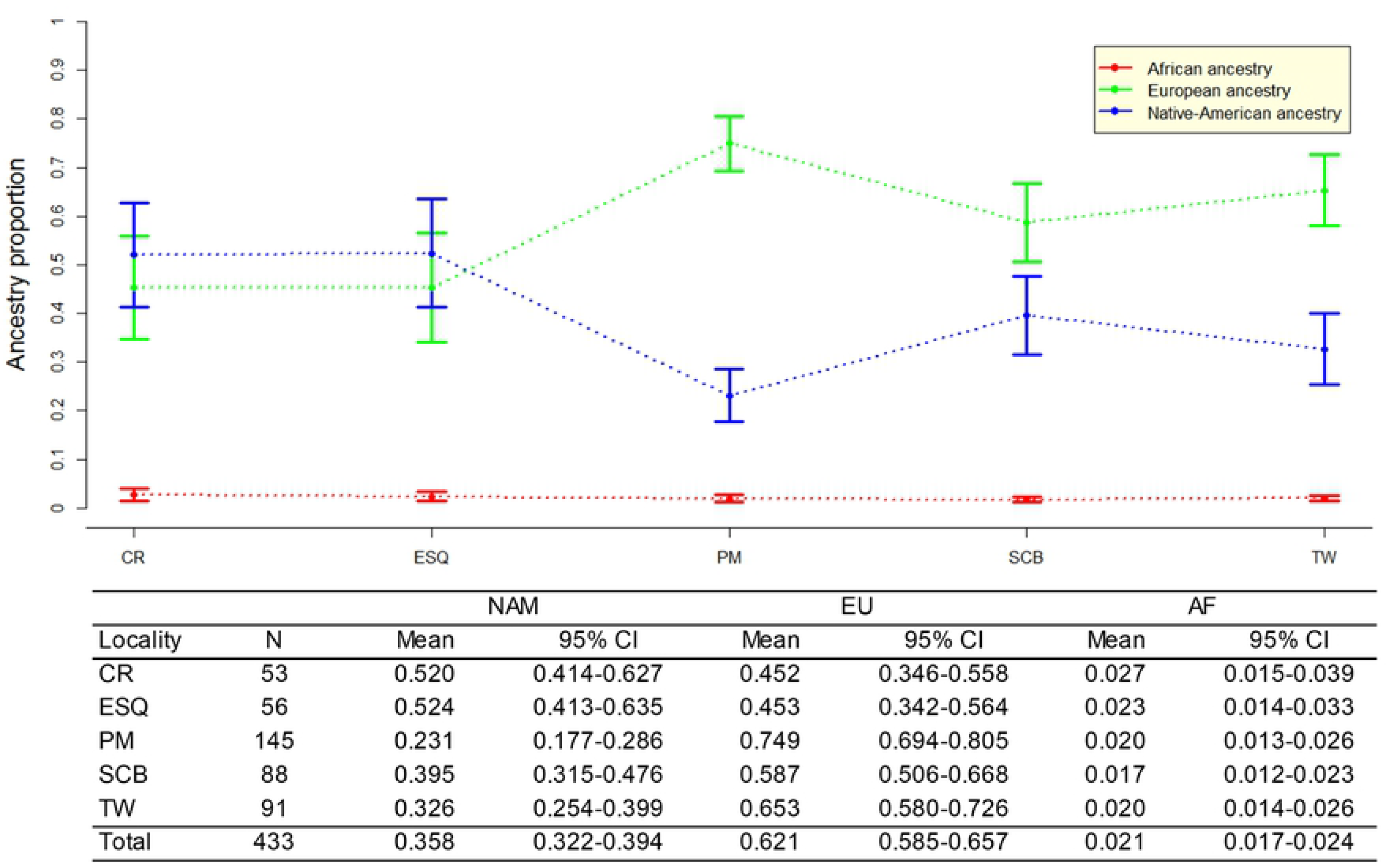
Average individual ancestry and the associated 95% CI for the five studied population. The blue line represents the Native American component (NAM), the green line the European component (EU), and the red line the African component (AF).

The Kruskal-Wallis test showed that the differences observed for the Native American and European ancestry distributions among the five samples are statistically significant (H ≥ 45.285; p<0.0001), while no significant differences (H = 4.631; p = 0.33) were observed for the African ancestry. In all pairwise comparisons, significant differences (p<0.05) were observed for both for Native American and European ancestry proportions, except between Comodoro Rivadavia and Esquel (NAM p = 0.389, EU p = 0.439) and between Trelew and San Carlos de Bariloche (NAM p = 0.1124, EU p = 0.152).

The overall results showed three main groups among central Patagonian localities: i) Comodoro Rivadavia and Esquel, ii) Trelew and San Carlos de Bariloche, and iii) Puerto Madryn.

As observed in our previous study based on autosomal Alu markers [18], the admixture process was not homogeneous across the Patagonia region, showing local differences on Native American and European ancestries. This heterogeneity is mainly related to different geographic origin of the inhabitants and their ancestors, different foundational histories and socio-economic process, which are widely discussed at the end of this section.

Aiming to compare the ancestry distribution between the three main regions of the country (North/Center/South), our data was analyzed together with those obtained by Toscanini et al. [15], for the same set of 46 AIM-Indels. As also found by Toscanini et al. [15], the Chi-square test showed that the ancestry varies between regions (p = 0.003). In this respect, the European contribution appeared to be markedly high in the Central region of Argentina (81.4%), while the Native American component increases toward the North (38.8%) and the South (37.8%) of the country (S1 Fig).

Differences on the ancestry of Argentinian regions were also found in other studies, based on 24 AIMs [12] and 99 AIMs [17] (S4 Table). For the Southern region, a high Native American contribution was registered in all studies. However, in Corach et al. [12], the European component appears to be overestimated in the South and in the North regions. This could be due to differences in the markers analyzed and/or in the populations sampled. In fact, as noted above, a large variation exists among South region populations, and it is possible that the same occurs in the North region. It is also worth noting that sampling in both, Corach et al. [12] and Toscanini et al. [15], was based on the place of residence of the individuals and no information on the birthplaces of the participants or their relatives is provided. The higher Native American contribution found in the South by Avena et al. [17], when compared to other studies, can also be explained by the heterogeneity of the populations studied from this region. Indeed, their study included samples from the two localities, Esquel and Comodoro Rivadavia, showing the highest Native American contributions, in this study and in previous studies using Alu autosomal markers (42.3% and 40.5%, respectively) [18], and mtDNA (78% and 73%, respectively) [3,5].

At regional level, the relatively high preservation of the Native American ancestry obtained in current urban Patagonia populations might be related to the late incorporation of this region to the National State (Law 1584, 1884), which allowed the Native populations to keep their autonomy for longer. Also, it might be related to the high migratory flow that the Patagonia region received from the inland provinces and from bordering countries, mainly Chile, with high Native American genetic background [4,36–38].

Likewise, at the local level, the admixture results obtained seem to be related to the birthplace of the participants and their ancestors, since a high variation was observed among grandparents’ birthplaces, for the five Patagonian samples. Comparing to the remaining samples, Esquel showed the lowest immigration from other regions, allowing for a better preservation of the original genetic background (Fig 4). Indeed, Esquel showed the highest average of Native American ancestry (52.4%; Fig 3), and the highest frequency of grandparents born in the same place (25.7%) (Fig 4).

**Fig 4.**
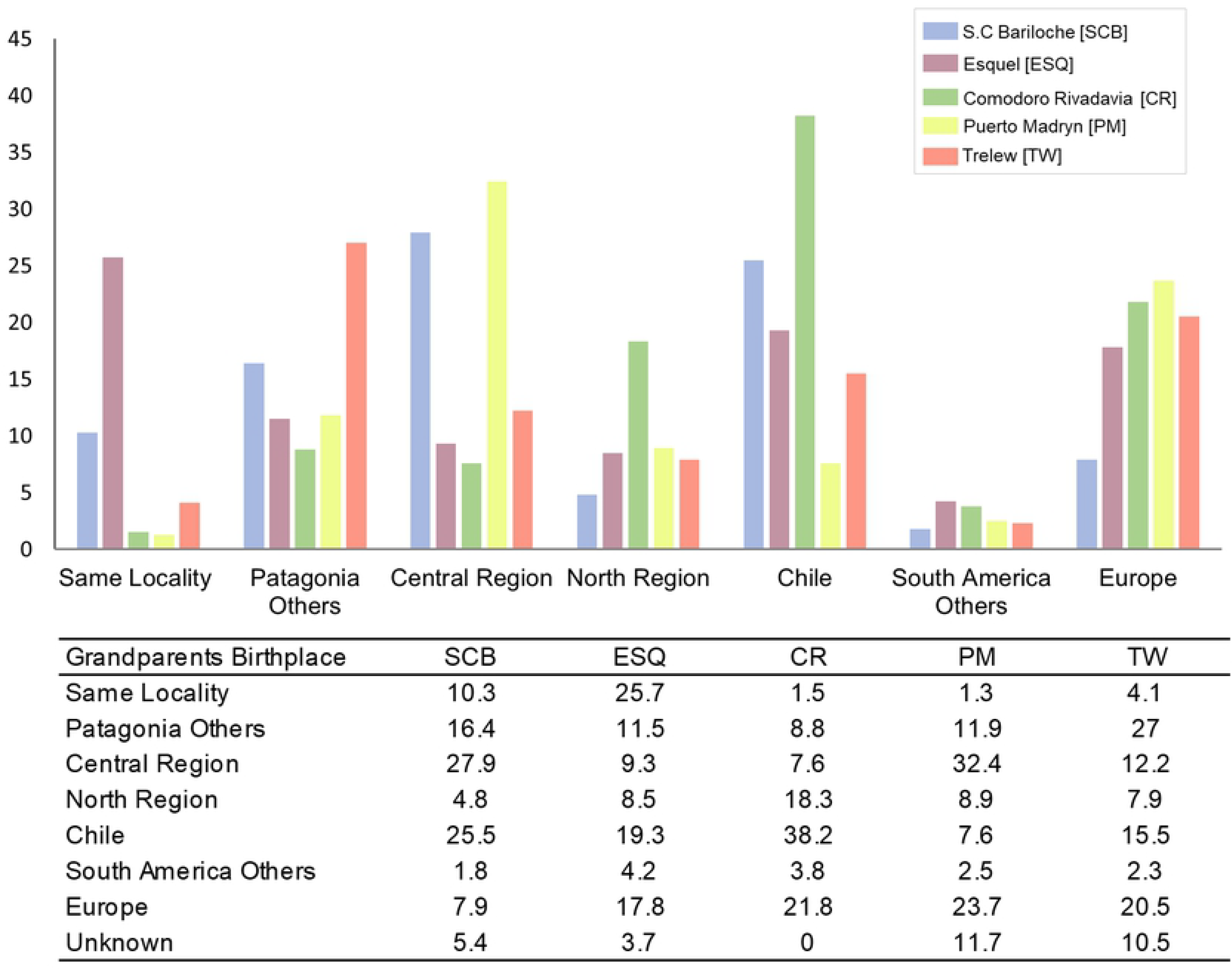
Proportion (%) of grandparents’ birthplace by locality.

This could be related with the gradual and diversified growth of the population, and a better preservation of the autochthonous origins [3,39]. Since before its foundation (1906) to the present day, Esquel has maintained a strong trade with Chile, being a zone of transport and gateway of migrants in both sides of the Andes (19.3% of the grandparents born in Chile). San Carlos de Bariloche, the most populated southern Andean city (512 inhabitants/km^2^), registered 39.5% of Native American ancestry (Fig 3), and 10.3% of the grandparents born in the same locality. However, 27.9% of the grandparents came from the Central region, mainly attracted by the tourism industry. As Esquel, San Carlos de Bariloche also has a strong migratory flow from Chile [4], as in the sample analyzed 25.5% of grandparents born in this neighboring country. Comodoro Rivadavia city was the second population with highest Native American ancestry (52%; Fig 3). It showed the highest proportion of grandparents from Chile (38.2%) and North West Argentina (18.3%) (Fig 4). Both immigrant groups, which preserve a high Native American ancestry [17,20,21,37,40], where mainly attracted to Comodoro Rivadavia by the exponential growth of the oil industry which started in 1907 and is still ongoing [5]. Meanwhile, Puerto Madryn showed the highest European ancestry (74.9%; Fig 3), and the highest proportion of grandparents from Europe (23.7%) and Central region (32.4%) (Fig 4), which presents the highest European ancestry of the country [12,15,17,21]. As mentioned in the introduction, between later 19th century and the end of the Second World War, a large number of Europeans arrived at the Central region of Argentina, who mainly settled in Buenos Aires province, and later spread to Pampean and littoral areas, attracted by the agriculture opportunities [1,2]. In Puerto Madryn, the major proportion of immigrants from the Central region is most likely related to the demand of qualified laborers for the aluminum industry. Puerto Madryn and Trelew, 65 km distant from each other, share the same foundational origin of Welsh, Spanish and Italian roots. Despite the common historical origin and geographic proximity, Trelew showed the highest proportion of grandparents born in other Patagonian localities (27%), likely attracted to Trelew by labor sources in the textile industry and public administration [8].

The five cities sampled here have less than 160 years of existence and relative low population densities (i.e., Esquel has 27.9 inhabitants/km^2^). Therefore, migrations have had an important impact in the growth of their populations.

### Origins and ancestry

The information about grandparents’ birthplace (region/country) was compared with the proportions of Native American, European and African ancestries of the participants, to investigate a possible association between them. For this analysis, 85 individuals with the 4 grandparents born in the same geographic place were selected. These samples were grouped according to the grandparents’ origin in five major regions: Europe, Chile, North Argentina, Central Argentina, and South Argentina.

Not surprisingly, the results showed that the number of individuals with grandparents from Europe was highly associated to the European ancestry in the population. The average of European ancestry among individuals with all grandparents from Europe was 96.8% (Fig 5). This is in agreement with previous observations by Avena et al. [17], suggesting that genealogical information, such as the number of grandparents from Europe, could be a strong predictor of genetic ancestry in samples from Argentina.

**Fig 5.**
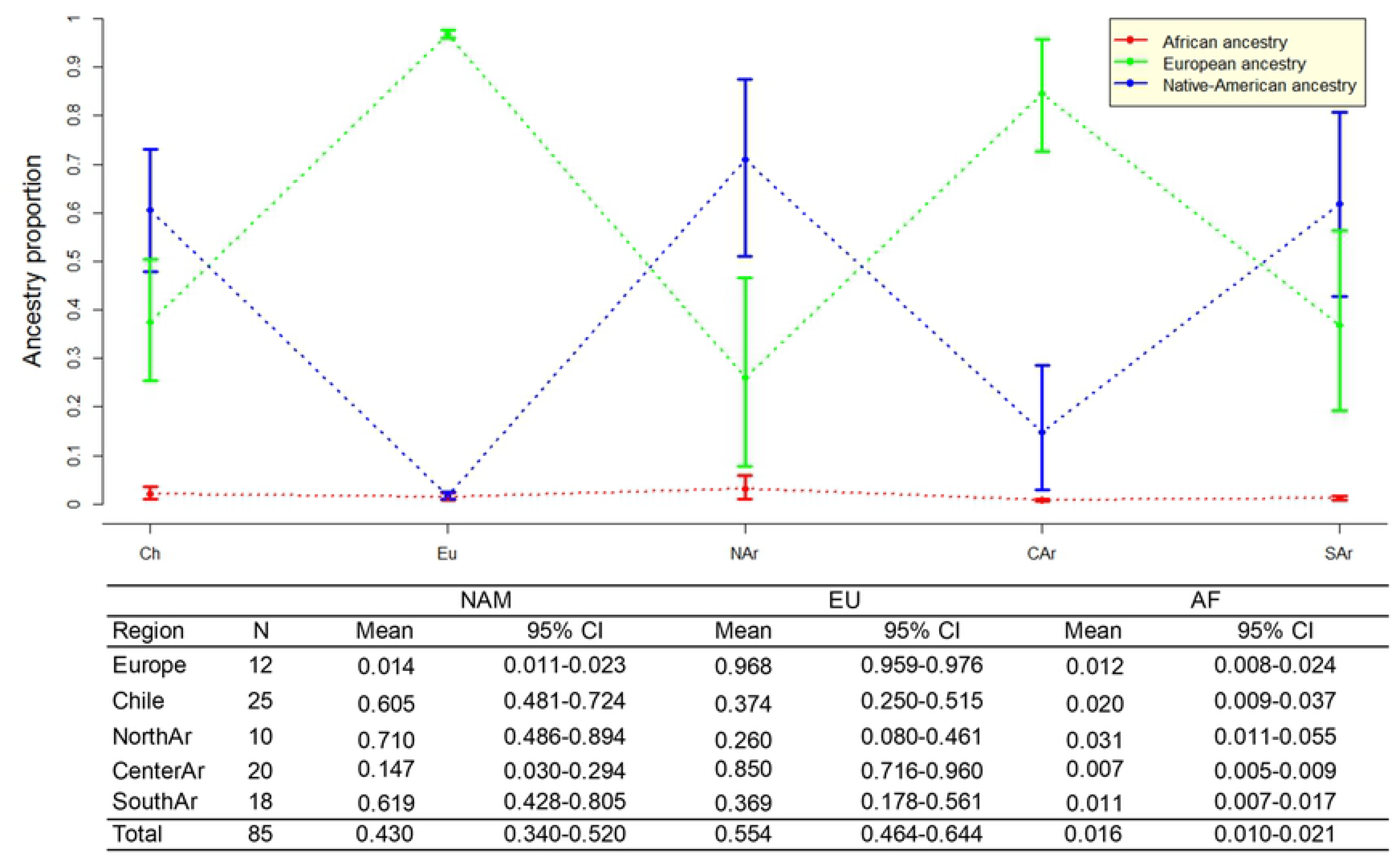
Average individual ancestry and the associated 95% CI obtained in samples with four grandparents born in the same geographic region. The blue line represents the Native American component (NAM), the green line the European component (EU) and the red line the African component (AF).

Also, a clear association was observed between the Native American ancestry and the number of grandparents born in Chile. The average Native American ancestry among individuals with four grandparents from Chile was 60.5%. These findings are in accordance with the relatively high prevalence of the Native American ancestry (range 46.9%-49.5%) reported in Chilean Patagonia [37].

Comparing the three main regions of Argentina, an association was also observed between the grandparents’ birthplace and the average European and Native American ancestries of the individuals. As expected, a high correlation between European ancestry and birthplaces in the Central region of Argentina was observed. The average European ancestry among individuals with all grandparents born in the Central region was 85% (Fig 5). As previously mentioned, the large number of European immigrants who arrived in Argentina between 1860 and 1950 mainly settled in Buenos Aires city and in the rest of the Pampean region [1,2].

In contrast, the Native American ancestry increased and the European decreased when the four grandparents were born in the North (NAM = 71%) or in the South (NAM = 61.9%) regions of the country (Fig 5). These results reflect the historical and demographic information described in previous section, and is consistent with the genetic studies carried out on biparental [12,15,17,18,20] and uniparental markers [3,5–8,40].

When Kruskal-Wallis test was assessed, significant differences (H ≥ 43.495; p<0.001) were observed for the Native American and European ancestry distributions among participants with four grandparents born in the three main regions of Argentina, Europe or Chile. However, no significant differences (H = 10.219; p = 0.045) were detected for the African component. In accordance with the ancestry frequencies obtained for each region (Fig 5), the *post-hoc* analysis showed no significant differences (p>0.05), for the Native American and European ancestries, between donors with four grandparents born in Europe and in Center Argentina (NAM p = 0.126, EU p = 0.288), neither between donors with four grandparents born in Chile, North and South Argentina (NAM p = 0.288-0.490, EU p = 0.231-0.476). These results are consistent with the high average European ancestry (≥ 85%) in the samples with four grandparents born in Center Argentina and in Europe. Also, similar average Native American ancestry (≥ 60%) were observed among participants with four grandparents born in Chile, North and South Argentina (Fig 5).

### Geospatial representation of the ancestry

The results obtained showed a clear variation in ancestry proportions mainly associated to the geographical origin of the samples. In order to obtain a spatial representation of this variation, we performed a geostatistical modelling based on the geographic coordinates for the birthplaces of the participants and their individual ancestry. For this representation, 392 individuals born in North, Central and South Argentina, and in Chile were included (41 samples from other countries or with incomplete birthplace information were excluded). The distribution of the individual genetic ancestry among the samples included in this analysis varied from 0.4 to 99% Native American, 0.6 to 99.2% European, and 0.3 to 23.4% African contribution.

In agreement with previous studies [12,15,17,18,20,21,37], the maps show the highest levels of Native American ancestry in individuals born in the North-Western region (NAM = 58.3%, 95% CI: 37.1-75.9%) and in Argentinian and Chilean Patagonia (NAM = 46.4%, 95% CI: 41.6-51.2%) (Fig 6A). As observed by Muzzio et al. [21] and Avena et al. [17], the European ancestry was higher in individuals born in the North East and Central Argentina (EU = 80.6%, 95% CI: 75.6-85.6%) (Fig 6B). Finally, the African ancestry is poorly represented across the country, with the highest frequency observed in northern Argentina (AF = 4.3%, 95% CI: 1.9-7.1%) (Fig 6C). This is also in agreement with previous studies carried out in this region [21,40].

**Fig 6.**
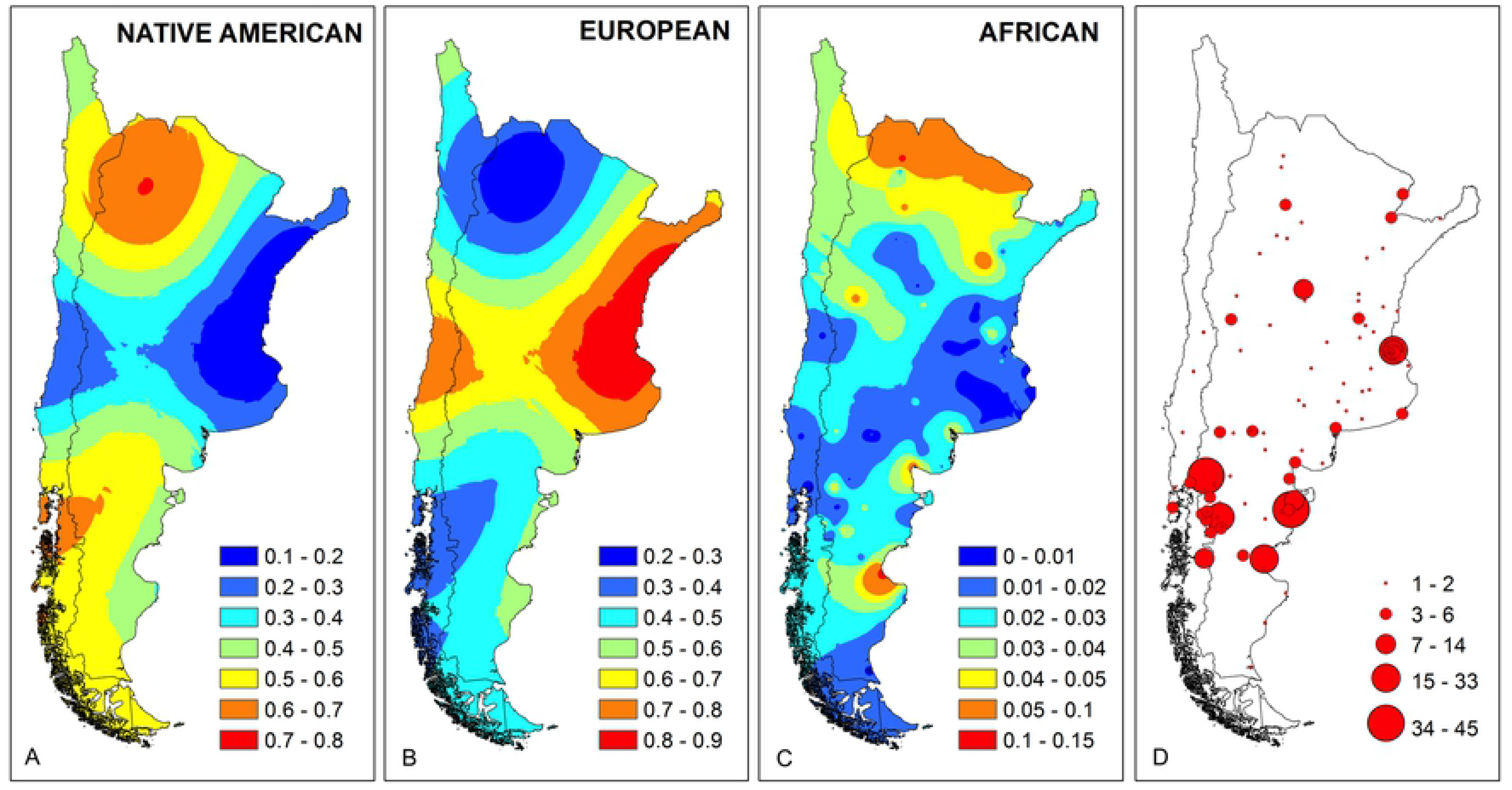
Maps of Argentina and Chile showing the distribution of Native American (A), European (B) and African (C) individual ancestry estimated in individuals who inhabit in Argentinian Patagonia. The birthplace of the individuals and the density are indicated by red dots (D).

### Genetic distance analysis

Using data from the 46 AIM-Indels, pairwise *F*_ST_ genetic distances were calculated between the five Patagonian populations from this study, and samples from six Argentinian provinces [15], six South American populations from Brazil [23], Bolivia [24] and Colombia [25,26], as well as the European, African and Native American HGDP-CEPH reference populations [22] (S5 Table).

Pairwise *F*_ST_ genetic distances showed significant differences (*p*-values < 0.000005) between the Argentinian populations and the parental and other South American populations except between Buenos Aires and South Brazil. Puerto Madryn showed low distances and no significant differences with Buenos Aires (*F*_ST_ = 0.0017, *p*-value = 0.1715) and La Pampa (*F*_ST_ = 0.0012, *p*-value = 0.2423). This is in agreement with the higher European ancestry registered in these populations, and with the birthplace of the grandparents of Puerto Madryn individuals, most born in the Central region of the country (Figs 4 and S1). On the other hand, no significant differences were observed between Esquel, Comodoro Rivadavia, Santa Cruz, and Tucumán (non-differentiation *p*-values > 0.0385; for a significance level of 0.00026, after applying the Bonferroni correction). This is consistent with the relatively high Native American ancestry present in these populations from the North and South regions (S1 Fig).

The pairwise *F*_ST_ matrix (S5 Table) was used to represent the genetic distances in a multidimensional-scaling plot (Fig 7).

**Fig 7.**
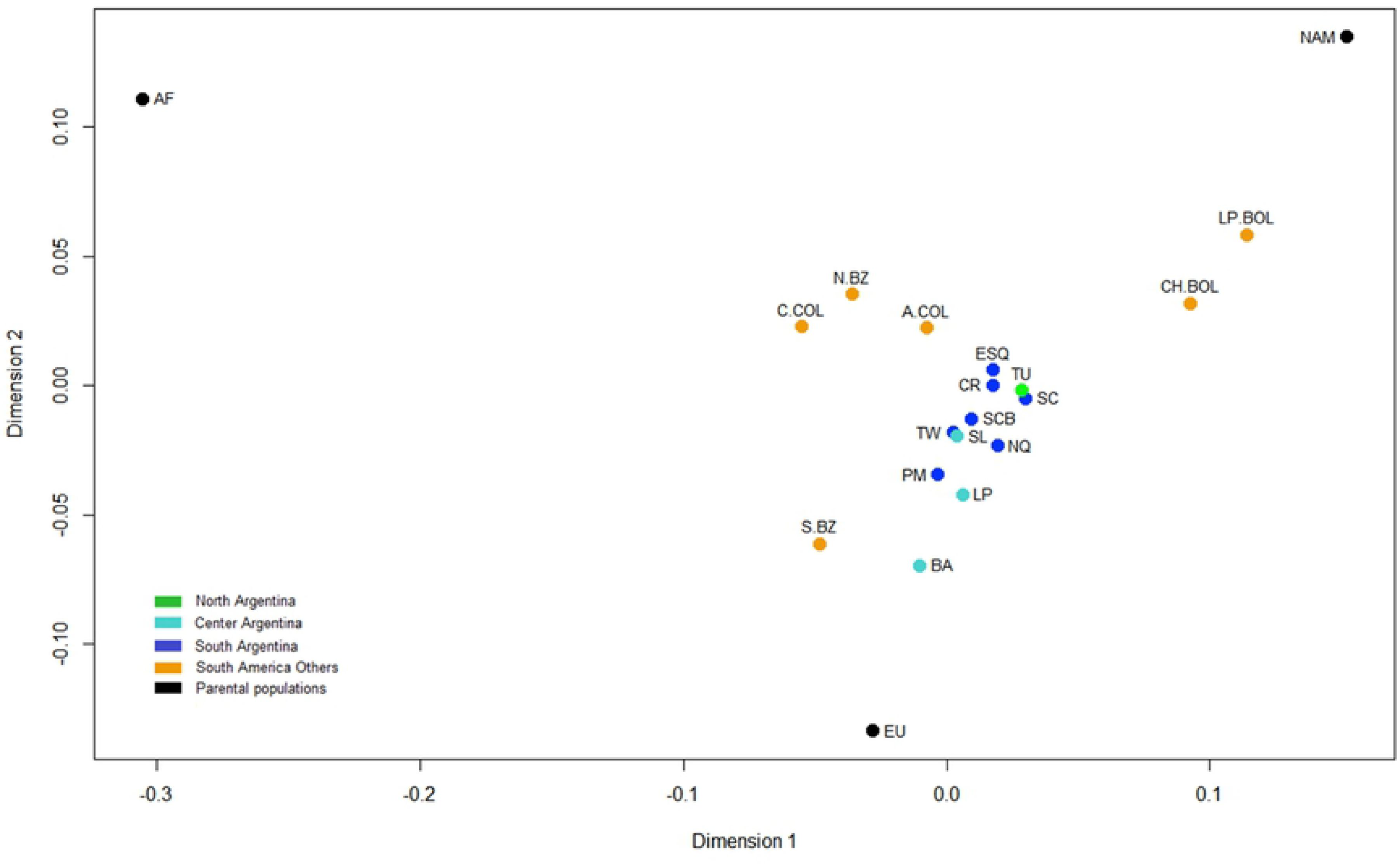
MDS plot of the *F*_ST_ pairwise genetic distances between eleven Argentine populations, other six South American populations and the three HGDP-CEPH parental populations. Stress = 0.10644. Population references are in S5 Table.

In accordance with the ancestry estimates, Argentinian populations exhibited an intermediate position between European and Native American reference groups. Puerto Madryn and Trelew are closer to Central region populations, while Esquel, San Carlos de Bariloche and Comodoro Rivadavia group together with other Patagonian populations and with Tucumán.

In summary, the location of Patagonian populations in the MDS plot closely resembles the results based on the ancestry estimates and are also in accordance with our previous genetic distances analysis based on uniparental and autosomal markers carried out in central Patagonia populations [6,7,18].

## Conclusion

This study constitutes the first analysis of the genetic ancestry in different populations of the Argentinian Patagonia region, complementing previous information obtained on uniparental and biparental markers. The appropriate representation of the studied samples, both in size and collection strategy, supports the obtained results. Likewise, the 46 AIMs analyzed prove to be robust and reliable for the estimation of population admixture, which has been supported by its strong association with genealogical information, mainly for European ancestry.

The different analyses performed allowed appreciating the intraregional differences in urban Argentinian Patagonia, even in a same province, where Puerto Madryn shows the highest European contribution, while Comodoro Rivadavia and Esquel preserve strong native roots. Nevertheless, as it was demonstrated in this study, the Native American contribution have different origins in each region and even in each locality, highlighting the importance of knowing the origin of the participants and their ancestors for the correct interpretation and contextualization of the genetic information.

Moreover, the differences observed among the five localities of the same region and of the same province, as in the case of Chubut, highlight the importance of taking into account the local historical demographic process in the analysis of the genetic composition of admixed populations. Accordingly, the admixture results, combined with the genealogical information, reveal that intra-regional variations are in agreement with the different geographic origin of the participants and their grandparents’, and with different local and social-demographic histories.

Finally, the heterogeneity among the populations analyzed emphasizes the need of additional studies involving rural populations to obtain an integral picture of Argentinian Patagonia, which is important for the correct interpretation of the genetic information in different studies, such as in medical, forensic and anthropological fields.

## Acknowledgments

The authors greatly thank to the participants, the Directors of the Hemotherapy Services (Laura Jones, María Del Carmen Ríos Part, Silvina Fleischer, Guillermo Manera, Claudia Tedeschi and Marcelo Furque), the hemotherapys’ technical teams for their support in collecting the samples, and Freya Chappell for carefully reviewing the writing.

## Supporting Information

**S1 Fig. Continental ancestry estimated for 11 Argentinian populations (5 from present work and 6 from Toscanini *et al.* [15], grouped in the three main regions of Argentina.**

(TIF)

**S1 Table. List of 46 AIM-Indels genotypes obtained in five urban Argentinian Patagonia populations**.

(XLS)

**S2 Table. Allele frequencies for 46 AIM-Indels in five urban Argentinian Patagonia populations**.

(XLS)

**S3 Table. Observed and expected values of heterozygotes and P-values for the exact test of Hardy-Weinberg equilibrium.**

(XLS)

**S4 Table**. **Average ancestry observed in the three main regions of Argentina for 24, 46 and 99 AIM sets.**

(XLS)

**S5 Table. Genetic distances (*F*ST) between 17 South American populations, and 3 parental populations (lower diagonal), and corresponding non-differentiation *P-* values (upper diagonal)**.

(XLS)

